# Ionic current correlations are ubiquitous across phyla

**DOI:** 10.1101/137133

**Authors:** Trinh Tran, Cagri T. Unal, Laszlo Zaborszky, Horacio Rotstein, Alfredo Kirkwood, Jorge Golowasch

## Abstract

Ionic currents, whether measured as conductance amplitude or as ion channel transcript levels, can vary many-fold within a population of identified neurons. This variability has been observed in multiple invertebrate neuronal types, but they do so in a coordinated manner such that their magnitudes are correlated. These conductance correlations are thought to reflect a tight homeostasis of cellular excitability that enhances the robustness and stability of neuronal activity over long stretches of time. Notably, although such ionic current correlations are well documented in invertebrates, they have not been reported in vertebrates. Here we demonstrate with two examples, identified mouse hippocampal granule cells and cholinergic basal forebrain neurons, that ionic current correlations is a ubiquitous phenomenon expressed by a number of species across phyla.

## Introduction

Ionic current levels in populations of identical neurons are extremely variable (Goldman, Golowasch, Marder, & Abbott, 2001; Leao, Li, Doiron, & Tzounopoulos, 2012; Liss et al., 2001; Olypher & Calabrese, 2007; Ransdell, Nair, & Schulz, 2012; Roffman, Norris, & Calabrese, 2012; Schulz, Goaillard, & Marder, 2006; Swensen & Bean, 2005). This poses the question of how neurons of a given type manage to generate consistent activity patterns despite the sometimes enormous variability of the currents they express. One mechanism that has been proposed is the co-regulated expression of ionic channels, which is revealed as correlations of conductances or transcript numbers in populations of identical cells (O’Leary, Williams, Caplan, & Marder, 2013; O’Leary, Williams, Franci, & Marder, 2014). The correlated expression of ionic currents, maximal conductances and ion channel transcript levels among populations of identical neurons have been observed in several neuronal cell types of invertebrate species (Khorkova & Golowasch, 2007; MacLean, Zhang, Johnson, & Harris-Warrick, 2003; Ransdell, Nair, & Schulz, 2013; Schulz, Goaillard, & Marder, 2007; Tobin, Cruz-Bermudez, Marder, & Schulz, 2009). However, that has typically been assumed to be an invertebrate idiosyncrasy. Evidence of their existence in vertebrates is largely anecdotal or indirect (McAnelly & Zakon, 2000; Swensen & Bean, 2005), and the only existing report of current correlations in mammals shows a correlation of voltage dependence or kinetic parameters (Amendola, Woodhouse, Martin-Eauclaire, & Goaillard, 2012; McAnelly & Zakon, 2000). Nevertheless, there is ample theoretical work that suggests that ionic current amplitude correlations allow neurons to express similar patterns of activity despite expressing widely different ionic current amplitudes by keeping relative levels of different current types constant (Hudson & Prinz, 2010; Lamb & Calabrese, 2013; O’Leary & Marder, 2016; Olypher & Calabrese, 2007; Rotstein, Olarinre, & Golowasch, 2016). Added to this, there is evidence that the expression of ionic current correlations is a highly regulated phenomenon (Khorkova & Golowasch, 2007), suggesting that correlations play important roles in the long-term dynamics of neuronal activity, in the regulation of the robustness of this activity, or both.

Here we test the hypothesis that ionic current correlations are widely distributed across animal species, and demonstrate that ionic current amplitude correlations are also expressed in mammalian neurons. We conclude that this is a ubiquitous phenomenon observed in species across phyla.

## Methods

We report observations from two different cell types: hippocampal granule cells (GC) from the upper blade of the dentate gyrus (DG) from 114-128 days (∼4 months) old male and female C57BL/6 mice (Jackson Laboratories, Bar Harbor Maine), and choline-acetyl transferase positive (ChAT^+^) cells from basal forebrain (BF) of 30-90 day-old BAC transgenic mice expressing enhanced green fluorescent protein (eGFP) under the promoter of the enzyme choline acetyltransferase, ChAT (B6.Cg-Tg(RP23-268L19- EGFP)2Mik/J, Jackson Laboratories, Bar Harbor Maine).

All experiments were performed in accordance with the U.S. Public Health Service Policy on Humane Care and Use of Laboratory Animals, the National Institutes of Health Guidelines for the Care and Use of Animals in Research, and approved by the Rutgers University Institutional Review Board and by the Institutional Animal Care and Use Committee at Johns Hopkins University.

### Hippocampus granule cells

C57BL/6 adult (8-10 weeks old) mice of both sexes were “entrained” for at least 2 weeks in light tight compartments with 12 hour dark/light cycles. For slice preparation, the mice were removed from their cages 15 minutes before the Light-to-Dark transition (scheduled at 10 AM). The mice were first deeply anesthetized with isofluorane and then perfused transcardially with cold dissection buffer (5 ml at 10 ml/min) containing 92 mM *N*-methyl-D-glucamine (NMDG), 2.5 mM KCl, 1.25 mM NaH_2_PO_4_, 30 mM NaHCO_3_, 20 mM HEPES, 25 mM glucose, 2 mM thiourea, 5 mM Na-ascorbate, 3 mM Na-pyruvate, 12 mM N-acetyl cysteine, 0.5 mM CaCl_2_ and 10 mM MgSO_4_ pH adjusted to 7.4. After decapitation brains were removed quickly, and acute hippocampal slices (300 μm) were made as described (Boric, Munoz, Gallagher, & Kirkwood, 2008) in ice-cold dissection buffer bubbled with a mixture of 5% CO_2_ and 95% O_2_. The slices were allowed to recover for 15 min at 30°C in dissection buffer and then for one hour at room temperature in artificial cerebrospinal fluid (ACSF): 124 mM NaCl, 5 mM KCl, 1.25 mM NaH_2_PO_4_, 26 mM

NaHCO_3_, 10 mM dextrose, 1.5 mM MgCl_2_, and 2.5 mM CaCl_2_ bubbled with a mixture of 5% CO_2_ and 95% O_2_.

All recordings were done in a submerged recording chamber superfused with ACSF (30 ± 0.5°C, 2 ml/min). Visualized whole-cell voltage-clamp recordings were made from GCs from the upper blade of the DG with borosilicate glass patch pipettes (3-6 MΩ) filled with intracellular solution containing the following: 130 mM K-gluconate, 10 mM KCl, 0.2 mM EGTA, 10 mM HEPES, 4 mM MgATP, 0.5 mM Na_3_GTP, 10 mM Na-phosphocreatine (pH 7.2–7.3, 280–290 mOsm). GC express a large number of ionic currents (Morgan & Soltesz, 2010; Santhakumar, Aradi, & Soltesz, 2005), but several can be measured without the need of chemical inhibitors. Membrane currents were recorded in the presence of 20 μM 6-cyano-7-nitroquinoxaline-2,3-dione (CNQX), 100 μM 2-amino-5- phosphonovaleric acid (APV) and 10 μM bicuculline methiodide (BMI) to block fast synaptic transmission. Series resistance was < 20 MΩ (range 6-20M), and average input resistance was 81.9 ± 21.7 MΩ (range: 64.4 to 115.0 MΩ) were studied. Series resistance compensation of at least 80% was always used and currents were corrected for series resistance-induce voltage errors. All drugs were purchased from Sigma Aldridge or R&D (Tocris).

### Basal forebrain (BF) cells

The B6.Cg-Tg(RP23-268L19-EGFP)2Mik/J mice were processed exactly as described in (Unal, Golowasch, & Zaborszky, 2012).

### Ionic currents and conductances

We measured the following currents: delayed rectifier K^+^ (*I*_*Kd*_), transient A-type K^+^ current (I_A_), inward rectifier K^+^ current (*I*_*Kir*_), a fast early inactivating transient inward current (which we label *I*_*in*_, see below), hyperpolarization-activated inward current (*I*_*h*_), and the linear leak current (*I*_*leak*_) (Fig. 1). GCs were clamped both at a holding voltage (V_h_) of either -40 and -90 mV and voltage steps of 500-600 msec duration were applied in 10 mV increments at 0.33 Hz to measure *I*_*Kd*_, *I*_*in*_, and *I*_*leak*_. For *I*_*Kir*_we applied 800 msec pulses from V_h_=-40 mV in -10 mV increments. We measured *I*_*Kd*_(Fig. 1c, d) after leak subtraction as the current at the end of a step to +40 mV from V_h_ = -40 mV in hippocampal GCs (in BF ChAT^+^ cells we measured *I*_*Kd*_at +10 mV). In each case K^+^ current conductances were calculated by dividing the current by the driving force using the calculated *E*_*K*_(-84 mV in hippocampal GCs, -99 mV in ChAT^+^ BF cells). In BF ChAT^+^ cells *I*_*A*_was measured by subtracting the currents obtained from V_h_ = -40 mV from those measured from V_h_ = -90 mV, and *g*_*A*_was calculated from currents measured at +10 mV. *I*_*Kir*_(GCs only, see Fig. 1a, b) and *I*_*h*_(BF ChAT^+^ cells only) was measured after leak subtraction at the end of a voltage step to -120 mV, and *g*_*Kir*_was calculated by dividing the current by the driving force with *E*_*Kir*_measured from the same cell’s current-voltage (I-V) curve (-70.5 ± 2.1 mV, Fig. 1b). *g*_*h*_was calculated assuming a reversal potential of -10mV. *I*_*in*_is the early transient inward current we observed in GCs (Fig. 1c) that peaks at around -20 mV (Fig. 1d) from either a V_h_ = -40 or -90 mV. We did not observe a noticeable difference in amplitude as a function of V_h_. We refer to this current as *I*_*in*_because we did not attempt to further characterize the current for this project. We believe that it is likely to be dominated by a high threshold Ca^++^ current, perhaps with a Na^+^ current contribution. Finally, *g*_*leak*_was calculated as the slope of the I-V curve between -70 and -40 mV (Fig. 1a, d), which is predominantly a linear component.

**Fig. 1.**
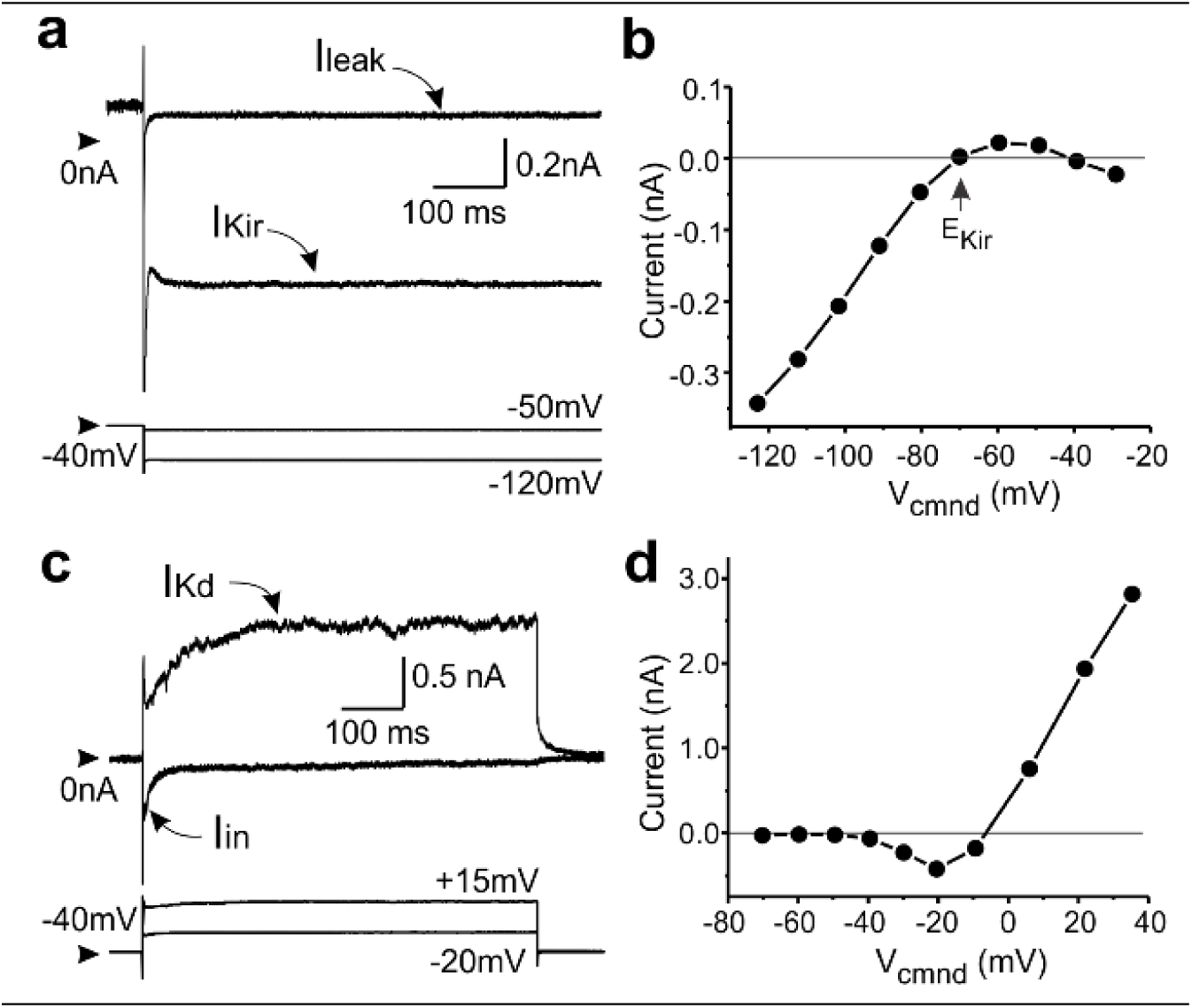
*Several ionic currents can simultaneously be measured in mouse hippocampal DG granule cells. I*_*Kd*_, *I*_*in*_, *I*_*Kir*_and *I*_*leak*_were measured with whole cell patch clamp in identified GCs. **a**. Inward rectifier (*I*_*Kir*_) and leak currents. **b**. Sample I-V curve for *I*_*Kir*_showing *E*_*Kir*_measurement. **C**. Slow delayed rectifier (*I*_*Kd*_) and early inward current (*I*_*in*_). **d**. Sample I-V curve measured at ∼20 msec from step onset. In a and c, top traces show currents; bottom traces show the pipette potentials at which the currents were measured. Synapses were blocked with APV, CNQX and bicuculline. Arrowheads show 0 nA (top traces) and -40 mV (bottom traces).

### Data analysis

Statistical analysis was performed using SigmaStat (Systat Software, Inc., San Jose, CA). Averages are represented as means ± SD and compared with t-Student tests for independent samples. Pearson product-moment correlation coefficients were calculated to reveal correlations between different variables. The Kolmogorov-Smirnov test was used to determine the normality of distributions.

## Results and Discussion

### Hippocampal granule cells

We recorded from 30 hippocampal GCs from the upper blade of the dentate gyrus (DG) from male (2) and female (3) mice at either the end of the “day” or the end of the “night” of 12h light-dark cycle entrained animals. Synaptic inputs were all blocked with APV, CNQX and bicuculline (Methods). We did not detect any significant differences between females and males and the data are thus pooled. To maximize the number of cells recorded we focused on four distinct ionic currents that can be studied without the need to add pharmacological agents (Fig. 1): the delayed rectifier (*I*_*Kd*_), an inward rectifier K^+^ current (*I*_*Kir*_), a fast transient inward current (*I*_*in*_), and a leak current (*I*_*leak*_).

**Figure 2.**
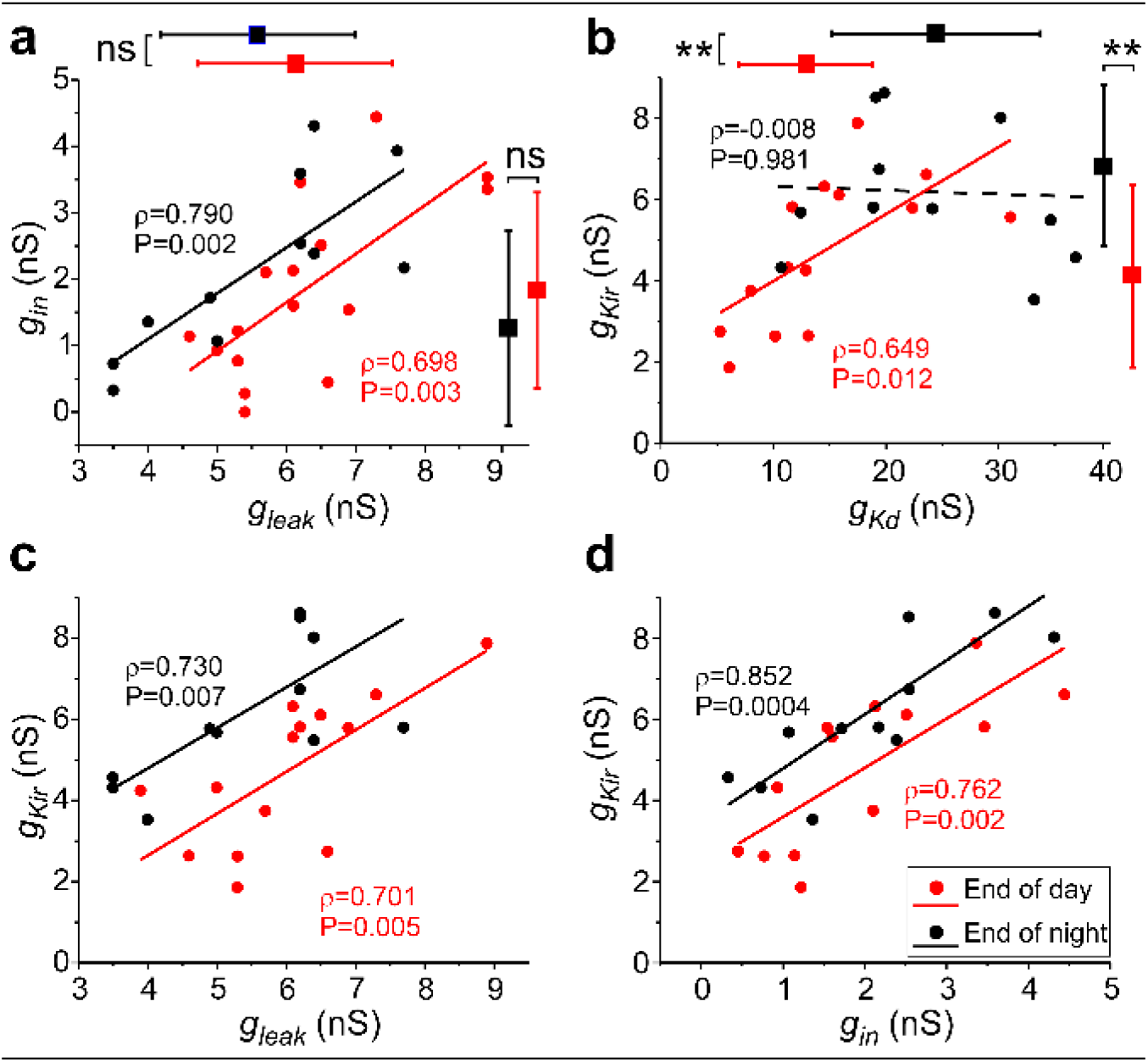
*Ionic conductance correlations in hippocampal GCs of 4 month-old mice. g*_*Kd*_, *g*_*in*_, *g*_*Kir*_and *g*_*leak*_are plotted against each other for data obtained at the end of the day (red) and end of night (blue)12h light-dark cycle). Only those pairs that showed significant correlations are shown. Pearson-moment correlation coefficients and their statistical significance, as well as regression lines, are shown in each panel. Means and SD bars for each conductance are shown to the right and top of the plots in a and b. ** P<0.001 (t-test).

As shown in figure 2 the variability of the conductance values of all currents measured both during the day and night was very large, with conductance ranges (ratio of max/min) as low as 2.2 for *g*_*leak*_(night) and as high as 15.9 for *g*_*in*_(day): 2.3 for *g*_*leak*_(day), 14.2 for *g*_*Kd*_(day), 4.2 for *g*_*Kir*_(day), 3.5 for *g*_*Kd*_(night), 2.8 for *g*_*Kir*_(night), and 13.1 for *g*_*in*_(night). Notably, all 4 conductances show statistically significant correlations (Pearson product-moment correlation, at P<0.05) in some subset of pairwise combinations (Fig. 2). The pairs *g*_*Kd*_*-g*_*leak*_and *g*_*Kd*_*-g*_*in*_did not show a significant correlation either during the day or night cycle, and are thus not plotted (Person-moment correlation coefficients and P values are shown in Figure 2). On the other hand, the pairs *g*_*leak*_-*g*_*in*_, *g*_*leak*_-*g*_*Kir*_, *g*_*in*_-*g*_*Kir*_, and *g*_*Kd*_-*g*_*Kir*_showed highly significant correlations even after adjusting for multiple comparisons (false discovery rate method (Curran-Everett, 2000)). There was one notable exception: the pair *g*_*Kd*_-*g*_*Kir*_showed no correlation during the night (ρ=-0.008, P=0.981) while the correlation during the day was strong and highly significant (ρ=-0.649, P=0.012). We compared these two slopes (slope end of night = -0.035, n=14, slope end of day = 2.555, n=12) using a Fisher z statistic (z(12,14) = 1.739) and found it to be significant at P=0.082. Although the normal cut off to consider a difference statistically significant is P=0.05, given that this value is essentially arbitrary, we take this result as a strong indication that there is a likely circadian regulation in the correlation of this pair, and only this pair, of conductances from among those we measured in this study. Importantly, the mean values of the two K^+^ conductances change between day and night (Fig. 2b side bars), while the mean values of *g*_*leak*_and *g*_*in*_do not significantly differ from each other (Fig. 2a, side bars).

Altogether the results indicate that ionic conductances in DG cells do vary in a correlated manner, and suggest a circadian-like control of some of these correlated pairs.

### Basal forebrain ChAT ^*+*^ neurons

We recorded from a total of 17 BF ChAT^+^ neurons. We analyzed three ionic currents, *I*_*A*_, *I*_*Kd*_and *I*_*h*_. *I*_*A*_and *I*_*Kd*_were expressed in all 17 cells (see example in Fig. 3a), but *I*_*h*_was measurable only in 10 of those cells. The range of conductances (ratio of max/min) was broad, similar to hippocampal GCs: 27.0 for *g*_*A*_, 14.0 for *g*_*Kd*_and 3.9 for *g*_*h*_. However, we detected a statistically significant correlation only for the *g*_*A*_-*g*_*Kd*_pair (Pearson product-moment correlation ρ=0.920, P=1.7x10^-7^) (Fig. 3b). For the remaining two pairs the correlations were characterized by: *g*_*A*_-*g*_*h*_pair (ρ=0.393, P=0.261); *g*_*Kd*_-*g*_*h*_pair (ρ=0.580, P=0.079).

**Figure 3.**
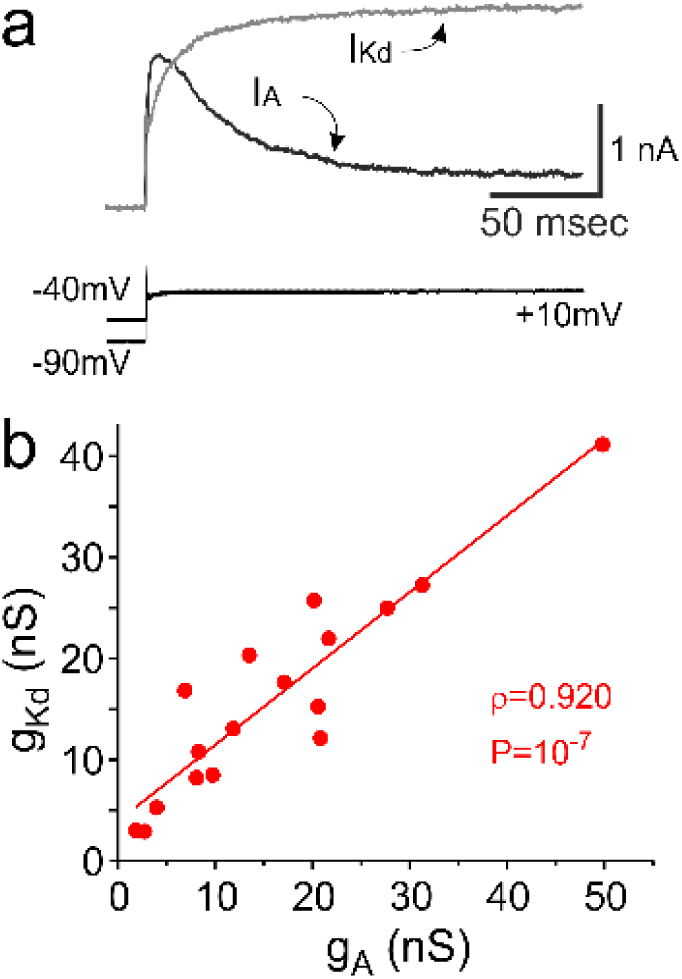
*Ionic currents and conductance correlations in adult mouse basal forebrain CHAT*^*+*^ *neurons.* **a.** Raw leak-subtracted *I*_*Kd*_(gray trace) and *I*_*A*_(black trace). Voltage steps used to elicit the currents are shown below the currents. **b**. *g*_*Kd*_and *g*_*A*_are plotted against each other. Pearson product-moment correlation coefficient, statistical significance, and regression line are shown.

Although the cell types sampled in this study are a minimal set of possible neuronal types, our results are entirely in line with the phenomenon thus far only characterized in invertebrate neurons, namely, that ionic current or conductances appear to be co-regulated in a very broad range of neurons also in vertebrates, specifically in mammals. Furthermore, we show that although not all currents appear to be correlated in any given cell type, a substantial fraction of them are. We predict that this will be shown in the future in an even wider range of neuronal subtypes in many different species. We think that these correlations reflect the existence of common regulatory pathways (see (O’Leary et al., 2013)) that are important in establishing cell type-specific setpoints in conductance space. These setpoints are not immutable, but can shift as neurons respond to persistent stimuli or factors, thus enabling these neurons to behave in cell-type characteristic ways while allowing the individual currents to vary in amplitude (cf. (Liu, Golowasch, Marder, & Abbott, 1998; O’Leary & Marder, 2016)). Like in invertebrates, multiple ionic current types appear to be correlated in mammalian neurons, which may be a cell type-specific characteristic (Schulz et al., 2007). Furthermore, we have shown evidence of what appears to be a regulatory mechanism of the correlations themselves in granule cells, which appears to be tied to a circadian mechanism. This would indicate that these correlations serve an important functional role that shifts between day and night. The cellular mechanism underlying this regulation is not known. However, consistent with this possibility, a regulation mechanism of ionic correlations has already been shown in crustacean neurons, in which the neuromodulatory environment determines the existence of correlations (Khorkova & Golowasch, 2007).

We conclude that ionic current amplitude correlations is a ubiquitous property of neurons across vertebrate as well as invertebrate species, and their functional significance remains to be determined in each case.

## Acknowledgements

This work was supported by NIMH grant R01_MH64711, NINDS grant R56_NS085330 (JG, HGR), NINDS grant R01_NS23945 (LZ).

## References

Amendola, J., Woodhouse, A., Martin-Eauclaire, M. F., & Goaillard, J. M. (2012). Ca(2)(+)/cAMP-sensitive covariation of I(A) and I(H) voltage dependences tunes rebound firing in dopaminergic neurons. The Journal of neuroscience: the official journal of the Society for Neuroscience, 32(6), 2166– 2181. doi:10.1523/JNEUROSCI.5297-11.2012 32/6/2166 [pii.

Boric, K., Munoz, P., Gallagher, M., & Kirkwood, A. (2008). Potential adaptive function for altered long-term potentiation mechanisms in aging hippocampus. The Journal of neuroscience: the official journal of the Society for Neuroscience, 28(32), 8034–8039. doi:10.1523/JNEUROSCI.2036- 08.200.

Curran-Everett, D. (2000). Multiple comparisons: philosophies and illustrations. Am J Physiol Regul Integr Comp Physiol, 279(1), R1–8.

Goldman, M. S., Golowasch, J., Marder, E., & Abbott, L. F. (2001). Global structure, robustness, and modulation of neuronal models. The Journal of neuroscience: the official journal of the Society for Neuroscience, 21(14), 5229–5238.

Hudson, A. E., & Prinz, A. A. (2010). Conductance ratios and cellular identity. PLoS Comput Biol, 6(7), e1000838. doi:10.1371/journal.pcbi.100083.

Khorkova, O., & Golowasch, J. (2007). Neuromodulators, not activity, control coordinated expression of ionic currents. The Journal of neuroscience: the official journal of the Society for Neuroscience, 27(32), 8709–8718. doi:27/32/8709 [pii] 10.1523/JNEUROSCI.1274-07.200.

Lamb, D. G., & Calabrese, R. L. (2013). Correlated conductance parameters in leech heart motor neurons contribute to motor pattern formation. PLoS One, 8(11), e79267. doi:10.1371/journal.pone.007926.

Leao, R. M., Li, S., Doiron, B., & Tzounopoulos, T. (2012). Diverse levels of an inwardly rectifying potassium conductance generate heterogeneous neuronal behavior in a population of dorsal cochlear nucleus pyramidal neurons. J Neurophysiol, 107(11), 3008–3019. doi:10.1152/jn.00660.2011 jn.00660.2011 [pii.

Liss, B., Franz, O., Sewing, S., Bruns, R., Neuhoff, H., & Roeper, J. (2001). Tuning pacemaker frequency of individual dopaminergic neurons by Kv4.3L and KChip3.1 transcription. EMBO J, 20(20), 5715– 5724.

Liu, Z., Golowasch, J., Marder, E., & Abbott, L. F. (1998). A model neuron with activity-dependent conductances regulated by multiple calcium sensors. The Journal of neuroscience: the official journal of the Society for Neuroscience, 18(2309-2320).

MacLean, J. N., Zhang, Y., Johnson, B. R., & Harris-Warrick, R. M. (2003). Activity-independent homeostasis in rhythmically active neurons. Neuron, 37(1), 109–120. doi:S0896627302011042 [pii.

McAnelly, M. L., & Zakon, H. H. (2000). Coregulation of voltage-dependent kinetics of Na(+) and K(+) currents in electric organ. The Journal of neuroscience: the official journal of the Society for Neuroscience, 20(9), 3408–3414.

Morgan, R. J., & Soltesz, I. (2010). Microcircuit model of the dentate gyrus in epilepsy. In V. C. e. al (Ed.), Hippocampal Microcircuits. NY: Springer

O’Leary, T., & Marder, E. (2016). Temperature-Robust Neural Function from Activity-Dependent Ion Channel Regulation. Curr Biol, 26(21), 2935–2941. doi:10.1016/j.cub.2016.08.06.

O’Leary, T., Williams, A. H., Caplan, J. S., & Marder, E. (2013). Correlations in ion channel expression emerge from homeostatic tuning rules. Proc Natl Acad Sci U S A, 110(28), E2645–2654. doi:10.1073/pnas.130996611.

O’Leary, T., Williams, A. H., Franci, A., & Marder, E. (2014). Cell types, network homeostasis, and pathological compensation from a biologically plausible ion channel expression model. Neuron, 82(4), 809–821. doi:10.1016/j.neuron.2014.04.00.

Olypher, A. V., & Calabrese, R. L. (2007). Using constraints on neuronal activity to reveal compensatory changes in neuronal parameters. J Neurophysiol, 98(6), 3749–3758. doi:00842.2007 [pii] 10.1152/jn.00842.200.

Ransdell, J. L., Nair, S. S., & Schulz, D. J. (2012). Rapid homeostatic plasticity of intrinsic excitability in a central pattern generator network stabilizes functional neural network output. The Journal of neuroscience: the official journal of the Society for Neuroscience, 32(28), 9649–9658. doi:10.1523/JNEUROSCI.1945-12.201.

Ransdell, J. L., Nair, S. S., & Schulz, D. J. (2013). Neurons within the same network independently achieve conserved output by differentially balancing variable conductance magnitudes. The Journal of neuroscience: the official journal of the Society for Neuroscience, 33(24), 9950–9956. doi:10.1523/JNEUROSCI.1095-13.201.

Roffman, R. C., Norris, B. J., & Calabrese, R. L. (2012). Animal-to-animal variability of connection strength in the leech heartbeat central pattern generator. J Neurophysiol, 107(6), 1681–1693. doi:10.1152/jn.00903.2011 jn.00903.2011 [pii.

Rotstein, H. G., Olarinre, M., & Golowasch, J. (2016). Dynamic compensation mechanism gives rise to period and duty-cycle level sets in oscillatory neuronal models. J Neurophysiol, 116(5), 2431–2452. doi:10.1152/jn.00357.201.

Santhakumar, V., Aradi, I., & Soltesz, I. (2005). Role of mossy fiber sprouting and mossy cell loss in hyperexcitability: a network model of the dentate gyrus incorporating cell types and axonal topography. J Neurophysiol, 93(1), 437–453. doi:10.1152/jn.00777.200.

Schulz, D. J., Goaillard, J. M., & Marder, E. (2006). Variable channel expression in identified single and electrically coupled neurons in different animals. Nat Neurosci, 9(3), 356–362. doi:nn1639 [pii] 10.1038/nn163.

Schulz, D. J., Goaillard, J. M., & Marder, E. E. (2007). Quantitative expression profiling of identified neurons reveals cell-specific constraints on highly variable levels of gene expression. Proc Natl Acad Sci U S A, 104(32), 13187–13191. doi:0705827104 [pii] 10.1073/pnas.070582710.

Swensen, A. M., & Bean, B. P. (2005). Robustness of burst firing in dissociated purkinje neurons with acute or long-term reductions in sodium conductance. The Journal of neuroscience: the official journal of the Society for Neuroscience, 25(14), 3509–3520. doi:10.1523/JNEUROSCI.3929- 04.200.

Tobin, A. E., Cruz-Bermudez, N. D., Marder, E., & Schulz, D. J. (2009). Correlations in ion channel mRNA in rhythmically active neurons. PLoS One, 4(8), e6742. doi:10.1371/journal.pone.000674.

Unal, C. T., Golowasch, J. P., & Zaborszky, L. (2012). Adult mouse basal forebrain harbors two distinct cholinergic populations defined by their electrophysiology. Front Behav Neurosci, 6, 21. doi:10.3389/fnbeh.2012.0002.

